# Functional and Optogenetic Approaches to Discovering Stable Subtype-Specific Circuit Mechanisms in Depression

**DOI:** 10.1101/498964

**Authors:** Logan Grosenick, Tracey C. Shi, Faith M. Gunning, Marc J. Dubin, Jonathan Downar, Conor Liston

## Abstract

**Background:** Using canonical correlation analysis (CCA), hierarchical clustering, and machine learning methods, we recently identified four subtypes of depression defined by distinct patterns of abnormal functional connectivity in depression-related brain networks, which in turn predicted differing clinical symptom profiles and individual differences in treatment response. However, whether and how dysfunction in specific circuits may give rise to specific depressive symptoms and behaviors remains unclear. Furthermore, this approach assumes that there are robust and stable canonical correlations between functional connectivity and depressive symptoms—an assumption that was not extensively tested in our earlier work.

**Methods:** First, we comprehensively re-evaluate the stability of canonical correlations between functional connectivity and symptoms, using optimized approaches for large-scale statistical testing, and we validate methods for improving stability. Next, we illustrate one approach to formulating hypotheses regarding subtype-specific circuit mechanisms driving depressive symptoms and behaviors and then testing them in animal models using optogenetic fMRI. We review recent work in this field and describe one example of this approach.

**Results:** Correlations between connectivity features and clinical symptoms are robustly significant, and CCA solutions tested repeatedly on held-out data generalize, but they are sensitive to data quality, preprocessing decisions, and clinical sample heterogeneity, which can reduce effect sizes. Generalization can be markedly improved by adding L2-regularization to CCA, which decreases variance, increases canonical correlations in left-out data, and stabilizes feature selection. This approach, in turn, can be used to identify candidate circuits for optogenetic interrogation in rodent models.

**Conclusions:** Multi-view approaches like CCA are a conceptually useful framework for discovering stable patient subtypes by synthesizing multiple clinical and functional measures. Optogenetic fMRI holds substantial promise for testing hypotheses regarding subtype-specific mechanisms driving specific symptoms and behaviors in depression.

---

Depression is not a unitary disease entity, but rather a heterogeneous neuropsychiatric syndrome that is thought to be caused by multiple distinct and interacting neurobiological mechanisms that may play unique roles in various patient subgroups (*1–6*). Pioneering earlier work in this field identified melancholic, atypical, seasonal, and other clinical subtypes of depression, each defined by sets of specific symptoms or other clinical characteristics that tend to co-occur in patient subgroups (*7–11*), but it has been challenging to identify subtype-specific neurobiological substrates that could be used as biomarkers. An alternative strategy for parsing heterogeneity would involve searching for patient subgroups defined by shared biological or objective cognitive and behavioral substrates, and then testing whether they predict clinical symptoms and outcomes—an approach that has already proven useful in psychosis, autism, and other neuropsychiatric disorders (*12–17*).

Motivated by these studies, our recent work identified four neurophysiological subtypes of depression defined by distinct patterns of altered functional connectivity as indexed by resting state fMRI in limbic and frontostriatal brain networks, which in turn predicted distinct clinical symptom profiles (*18*). In this work, we used canonical correlation analysis (CCA) to identify linear combinations of resting state functional connectivity (RSFC) features that predicted linear combinations of clinical symptoms, both of which could be used for either defining patient subtypes or for rating individual patients along continuous dimensions that capture unique aspects of brain dysfunction, consistent with multiple previous studies identifying correlations between RSFC features, clinical symptoms, and diagnostic status in depression (*19–26*).

However, while this approach was able to make useful predictions about clinical symptom presentations and treatment response probabilities, it also raised important questions. In identifying complex patterns of functional connectivity involving dozens of brain circuits associated with specific symptom combinations, it raised questions such as: which connectivity alterations in what circuits subserve particular symptoms and behavior; which are merely correlated with them; and how do connectivity alterations in multiple circuits interact? Such questions are difficult to answer using fMRI alone, but animal models hold promise for addressing them. Over the last ten years, new optogenetic approaches for experimentally manipulating brain circuit function in genetically defined cell types and topologically defined projections have begun to define causal relationships between circuit function and behavior (*27–32*), with important implications for both neurological (*33–35*) and psychiatric disease states (*30, 36–42*). Importantly, these methods can also be integrated with functional MRI and other noninvasive neuroimaging techniques that are widely used in humans, offering new opportunities for testing hypotheses and predictions derived from human neuroimaging studies (*36, 43*).

Here, we illustrate one such approach to formulating hypotheses regarding subtype-specific circuit mechanisms driving depressive symptoms and behaviors and then testing them in animal models using optogenetic fMRI. We review recent work in this field and describe one example of this approach, integrating results from our recent subtyping work with published optogenetic fMRI studies. Ideally, this approach could be extended to other datasets and samples that are clinically characterized but lack resting state fMRI data. But such extension would require a stable, low-dimensional embedding that could be readily and reliably generalized to new data—and such stability was not tested in our previous work. Relatedly, a recent preprint reported that CCA involving high-dimensional neuroimaging data tends to overfit, raising questions about the central assumption that functional connectivity alterations in depression-related brain networks are robust and reliable predictors of symptoms and behavior (*44*).

Therefore, we began by comprehensively re-evaluating whether functional connectivity alterations in depression are stable predictors of clinical symptoms using optimized approaches for large-scale statistical testing. We find that correlations between RSFC features and clinical symptoms are robustly significant, and that CCA solutions tested repeatedly on held-out data generalize well, but tend to overfit with increasing numbers of features. To overcome this obstacle, we show that this generalization can be markedly improved by adding L2-regularization to CCA, which decreases variance, increases canonical correlations in left-out data, and stabilizes feature selection. By testing which RSFC features appear most frequently across the best models, we can identify candidate circuits for optogenetic interrogation in rodent models. We then explain why multi-view approaches like CCA are a conceptually useful framework for discovering stable patient subtypes by synthesizing multiple clinical and functional measures, and can serve as a bridge for extending these results to new subjects across modalities. We conclude by reviewing recent advances in optogenetic fMRI and illustrating how this method could be used to test for subtype-specific circuit mechanisms driving particular depression-related behaviors.

## Methods and Materials

### Subjects

The analyses reported in Figs. 1–3 were designed to re-evaluate our approach in Drysdale et al., using state-of-the-art statistical methods to test whether depression-related RSFC alterations are statistically significant and stable predictors of clinical symptoms. Therefore, these analyses were conducted in the same “subtype-discovery sample” used in the Drysdale report, which comprised N = 220 subjects meeting DSM-IV criteria for a diagnosis of (unipolar) major depressive disorder and currently experiencing an active, non-psychotic major depressive episode at the time of the fMRI scan. All patients in this sample also met criteria for treatment resistance, with a history of failing to respond to at least two antidepressant trials of adequate dose and duration during the current episode. The subjects were recruited from outpatient clinics at Cornell (mean age = 42.1 years, 58.3% female) and the University of Toronto (mean age = 40.4 years, 57.3% female), using identical inclusion and exclusion criteria. Subjects were eligible for inclusion if they presented while seeking treatment for an active unipolar major depressive episode with a history of treatment resistance as defined above. Exclusion criteria were as follows: currently active substance use disorder; bipolar depression; a psychotic disorder; unstable medical conditions; a history of seizures or head injury with loss of consciousness; current pregnancy; and other contraindications for MRI (e.g. claustrophobia, implanted intracranial devices or cardiac pacemakers). Diagnoses were established by a trained clinician using a structured clinical interview (MINI or SCID). Subjects were not excluded on the basis of other psychiatric comorbidities or psychiatric medication usage. See Table 1 for details on medication status and psychiatric comorbidities. Symptom severity scores were quantified using the 17-item Hamilton Rating Scale for Depression (HAMD). As expected for a treatment-resistant population, they presented with moderate to severe total symptom scores (mean HAMD for Cornell subjects = 19.3; mean HAMD for Toronto subjects = 20.4).

**Figure 1:**
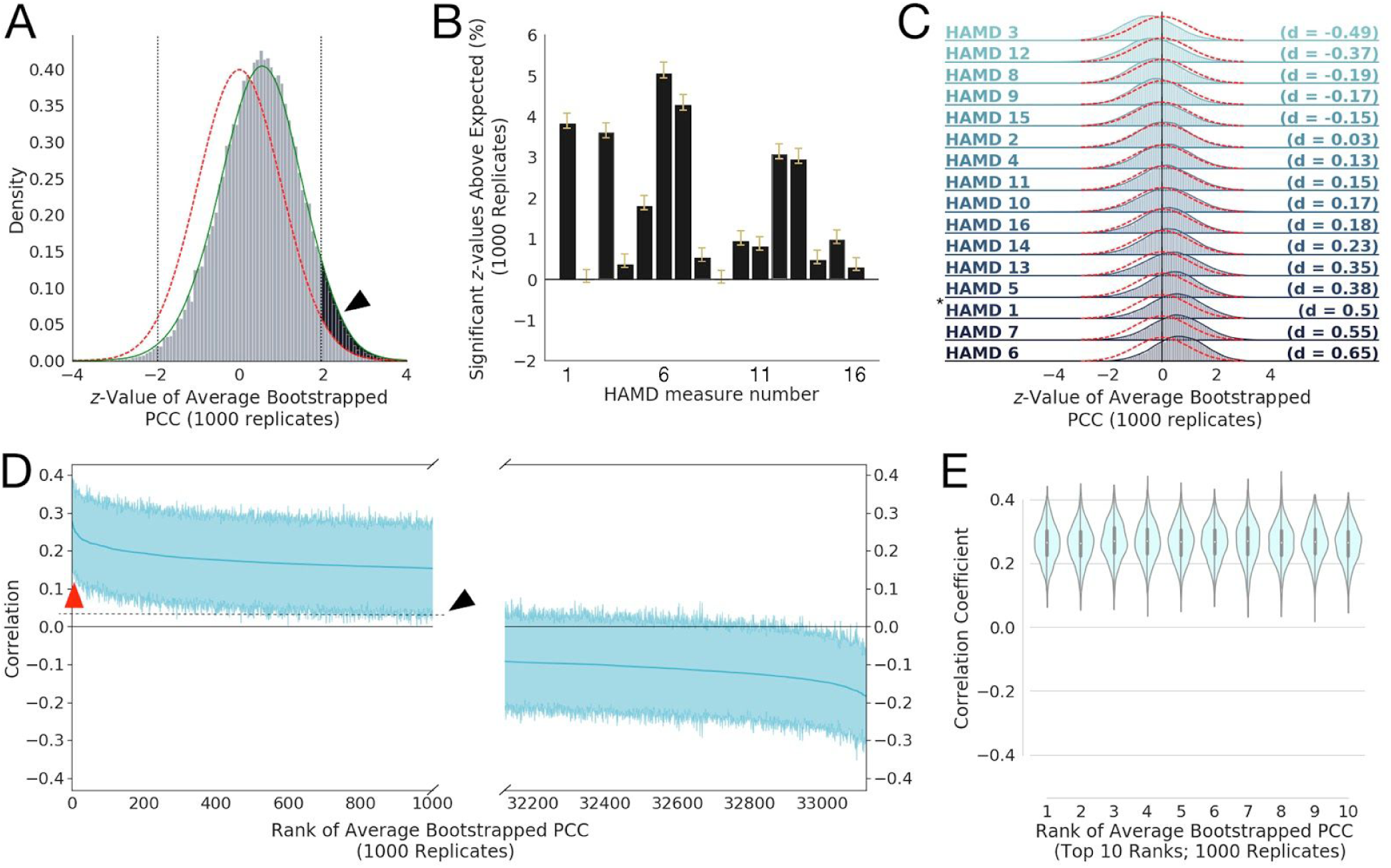
Robust correlations exist between Resting State Connectivity Features (RSFCs) and clinical measures (HAMD). **A)** Histogram of *z*-values for the 33,123 average Pearson Correlation Coefficients (PCCs) between Resting-State Functional Connectivity (RSFC) features and HAMD clinical measures (averaged over 1000 bootstrap replicates), compared to a standard 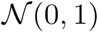 Gaussian distribution (red line) and with a smoothed kernel density estimate plotted over the histogram (green line; see Methods). The black arrow and shaded region show the area of the *z*-values that exceeds the area expected for a standard normal distribution (outside the two-sided significance criterion of *z* > ±1.96 for *p* < 0.05; shown as vertical dotted black lines). Note the empirical distribution of *z*-values has sample mean and standard deviation 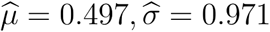, respectively, with the lower-than-expected sample variance resulting from correlations among the statistics; we correct for the effects of such inter-statistic correlations using the procedure in (*48*)(see Methods). **B)** Bar plots of the mean percentage of *z*-values that exceeded that expected by chance (e.g., the percentage above 2.5%, shown as the shaded black area in **A** for HAMD 1) for 1000 bootstrap replicates. Yellow whiskers on the bars denote 95% confidence intervals (corrected for multiple comparisons and data correlation using Bonferroni-Holm and (*48*), respectively). We see that HAMD measures 1, 3, 5, 6, 7, 12, and 13 have mean significant percentages well in excess of 1% more than expected under the null hypothesis. **C)** Histograms of *z*-values like that shown in **A** for all 16 HAMD clinical measures considered, ordered by effect size (Cohen’s d, given at right of each plot; magnitudes between 0.2 and 0.5 are considered small to medium effect sizes, between 0.5 and 0.8 are considered medium to large effect sizes). Red dotted lines denote the standard normal distribution. Asterisk (*) marks the distribution for HAMD1 shown in **A**. **D)** Bootstrapped PCCs for HAMD measure 1 for the 1000 most positive (left) and 1000 most negative (right) RSFC features (shaded regions shows 95% percentile-bootstrap confidence interval for the mean), ordered by mean correlation (thick blue line). Red arrow points to top 10 most positive-ranked RSFC features (shown in **E**); note both have confidence intervals excluding zero, indicating that while they cover an appreciable range, they are significantly different than zero across the 1000 bootstrap replicates and thus somewhat stable across bootstrap replicates. The black arrow and dotted line show the upward shift resulting from the positive shift of the distribution shown by the black arrow in panel **A**. **E)** Violin plot (with superimposed boxplots showing 1st and 3rd quartiles as black bar and the median as white point) of the top ten positive ranked RSFCs by average PCC to HAMD measure 1 (corresponding to red arrow in **D**), with mean 95% confidence intervals ± SD of [0.148 ± 0.0152, 0.376 ± 0.0119]. Note these look very similar, suggesting the rank order could easily change across replicates.

**Figure 2:**
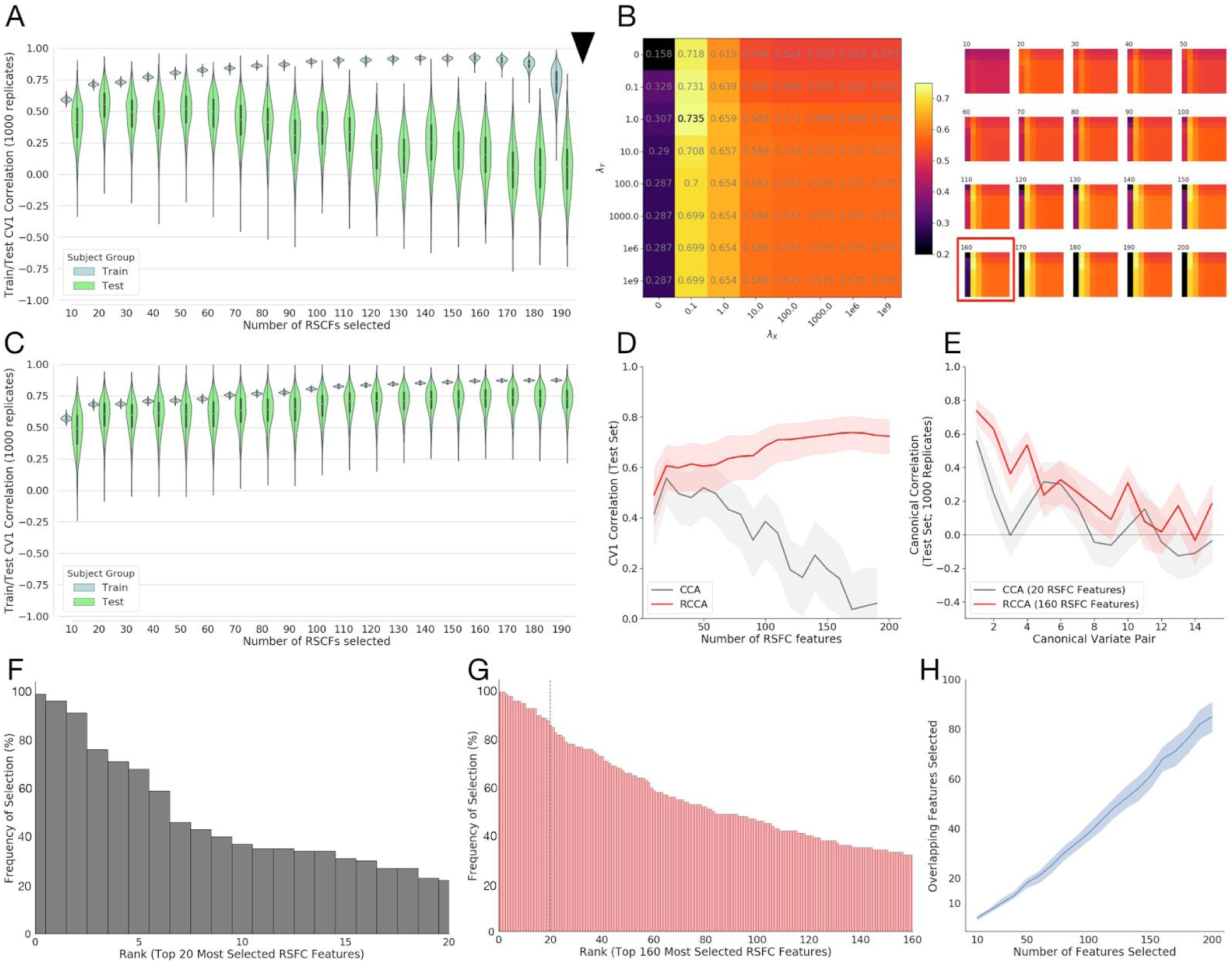
Stable train/test canonical correlations between RSFC features and clinical measures are improved by regularization. **A)** Violin plots (with superimposed boxplots) of correlations between the first canonical variates of a standard Canonical Correlation Analysis (CCA) on training data (90% of subjects) and test data (10% of subjects) for a range of features (10 to 190 by increments of 10) selected using the correlation method proposed in (*18*), with this procedure bootstrapped 1000 times for each number of features to yield the plotted distributions. Feature selection and CCA fitting was done on training data, separately for each bootstrap replicate, and then estimated CCA coefficients applied to the selected features in the held-out validation set to obtain test correlations. Test correlations for CCA peak at 20 features selected. Black arrow: standard CCA cannot be fit to more correlations than there are observations (in this case 90% of n=220, or 198 subjects). **B)** Median test rates fit over a grid of regularization parameters *λ_X_*, *λ_Y_* for each number of features selected. (Left) The grid corresponding to the best test correlations corresponding to using 160 RSFC features. The color of each square in the grid corresponds to the median test correlation (also printed in grey in the center of each square; colorbar at right gives hue values). (Right) Similar grids for other numbers of RSFC features (number of features selected shown above grid, test correlations shown in color only, not text). The best fit (160 features; shown on the left) is boxed in red box in the full set of fits on the right. Fitting more than 198 coefficients is possible. **C)** Violin plots (with superimposed boxplots) of correlations between the first canonical variates of the Regularized Canonical Correlation Analysis (RCCA) with the best regularization parameters (*λ_X_ =* 0.1, *λ_Y_* = 1, *N_F_* = 160) on training data (90% of subjects) and test data (10% of subjects) for the various numbers of features selected using the correlation method proposed in (*18*) (resampled 1000 times), as in **A**. Fitting more than 198 coefficients is possible. **D)** Test rates for the first canonical variate (CV1) as a function of the number of features selected for CCA (grey) and RCCA (red); shaded region shows 1st through 3rd quartile for the replicate fits. **E)** Test correlations between canonical variates 1–15 for the best fit from **A** (CCA fit in grey; 20 features), and the best fit from **C** (RCCA fit in red; 160 features); shaded region shows 1st through 3rd quartile for the replicate fits. **F)** Ordered (by top rank) histogram of the top 20 features chosen by the feature selection approach (from (*18*)) showing the percentage of times they were chosen across the 1000 subsampled replicate data sets. Just 3 features are selected more than 80% of the time. **G)** Ordered (by top rank) histogram of the top 160 features chosen by the feature selection approach showing the percentage of times they were chosen across 1000 subsampled replicates. 25 features appearing more than 80% of the time, dotted line denotes top 20 features; compare with **F**. **H)** Number of overlapping features in all pairwise combinations of 100 randomly chosen replicates as a function of number of features selected (dark blue line shows median and shaded region 1st through 3rd quartile across replicates). The median number of overlapping features selected increases approximately linearly with the total number of features selected.

**Figure 3.**
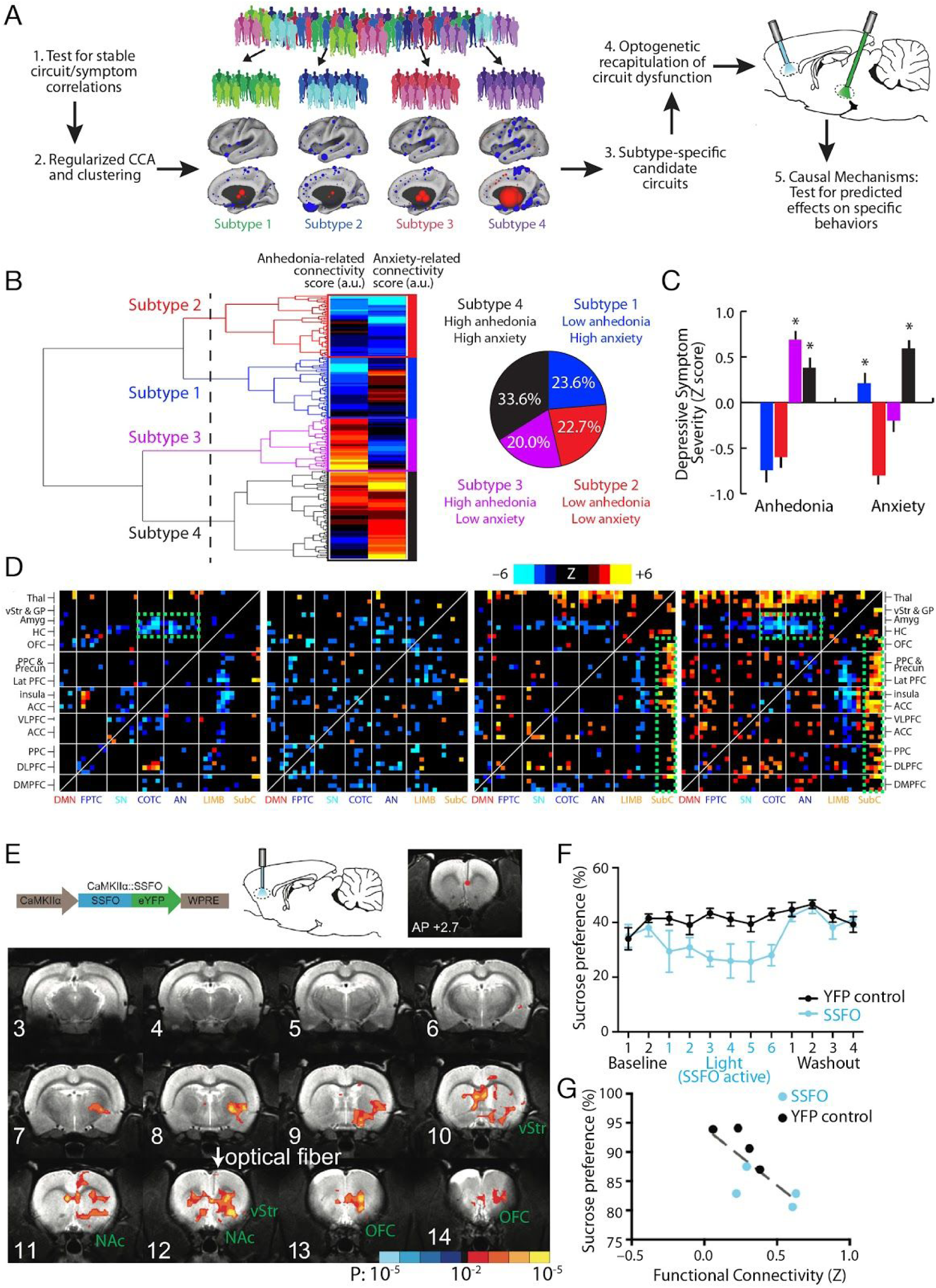
Optogenetic fMRI for interrogating subtype-specific circuit mechanisms in depression. **A)** Schematic illustration of a model for formulating hypotheses regarding subtype-specific mechanisms driving depressive symptoms and behaviors, and testing them in animal models using optogenetic fMRI. By first testing for robust and stable RSFC-clinical symptom correlations as in Fig. 1 and then using CCA and hierarchical clustering, relatively homogeneous subgroups of a heterogeneous MDD sample can be identified. These subgroups can be used to identify subtype-specific candidate circuits (see main text), and ofMRI can be used to test hypotheses about dysfunction in specific circuits driving specific behaviors, while also validating whether the RSFC effects evoked by the optogenetic manipulation resemble those observed in human subjects. **B)** In Ref. (*18*), hierarchical clustering on two canonical variates representing anhedonia- and anxiety-related RSFC revealed at least four clusters of patients in these two dimensions. The height of each linkage in the dendrogram represents the distance between the clusters joined by that link. The dashed line denotes 20 times the mean distance between pairs of subjects within a cluster. **C)** The four subtypes predicted significant group differences in anhedonia and anxiety (P < 0.005, Kruskal Wallis ANOVA) as indexed by item-level responses on the HAMD (item 7 and 11, respectively). Symptom severities are Z-scored with respect to the mean and standard deviation of all patients in the sample. Error bars = S.E.M. **D)** Heatmaps depicting subtype-specific patterns of altered functional connectivity for the top 50 neuroanatomical ROIs with the most subtype-specific RSFC features by Kruskal Wallis ANOVA. The color scale represents Wilcoxon rank sum test scores for the difference between patients in each subtype and matched healthy controls. The green boxes denote RSFC features discussed in the main text. For additional details on panels B-D, see Ref. (*18*). **E)** In Ref. (*36*), a viral vector (AAV/CaMKIIa/SSFO) driving SSFO expression in projection neurons was injected into mPFC, and an optical fiber implanted over the mPFC target was used to activate (blue light) and inactivate (amber light) the opsin during alternating rsfMRI scanning periods (300 s per scan). SSFO activation induced a pattern of increased functional connectivity between an mPFC seed (denoted by the red dot) and a network of structures depicted here, where colors denote the Z statistic (and associated P value) for RSFC changes in the opsin-on vs. opsin-off conditions (N = 4 rats, 14 runs). NAc = nucleus accumbens; OFC = orbitofrontal cortex; vStr = ventral striatum. **F)** Subjects (N = 8 SSFO rats, blue; N = 10 control rats, black) were assessed on the sucrose preference test during a 2-day baseline period, followed by 6 days with SSFO activated, followed by a 4-day “washout” period with SSFO off. SSFO activation reduced sucrose preference behavior (F(11,176) = 2.56, P = 0.0051, two-way repeated measures ANOVA), compared to subjects expressing a YFP control construct. **G)** Individual differences in RSFC between the mPFC seed and the ventral striatum correlated with sucrose preference behavior (R^2^ = 0.56, P = 0.03). Panels B-D and E-G were adapted from Refs. (*18*) and (*36*), respectively. See corresponding references for additional details.

**Table 1.** Subject demographics, medication status and psychiatric comorbidities. The analyses in Figs. 1–3 were implemented in the same dataset used in Ref. (*18*). Subjects recruited at the Toronto and Cornell sites were matched for age (p = 0.41), sex (p = 0.87), and depression severity (HAMD17 total score, p = 0.11). *Psychiatric medications listed as “Other” included benzodiazepines, non-benzodiazepine sedative-hypnotics, stimulants, and thyroid hormone. **Psychiatric comorbidities listed as “Other” included obsessive compulsive disorder, attention-deficit/hyperactivity disorder, Asperger Syndrome, and Tourette’s Syndrome.

|  | **Toronto Sample** | **Cornell Sample** |
| --- | --- | --- |
| **Number of Subjects** | 124 | 96 |
| **Age (mean)** | 40.4 years | 42.1 years |
| **Sex** | 57.3% female | 58.3% female |
| **HAMD17 Total Score (mean)** | 20.4 | 19.3 |
| **Psychiatric Medications** |  |  |
| **Antidepressant** | 59.7% | 57.3% |
| **Mood Stabilizer** | 16.9% | 17.7% |
| **Antipsychotic** | 17.7% | 15.6% |
| **Other\*** | 45.2% | 42.7% |
| **Psychiatric Comorbidities** |  |  |
| **Generalized Anxiety Disorder** | 4.8% | 5.2% |
| **Post-traumatic Stress Disorder** | 6.5% | 4.2% |
| **Social Anxiety Disorder** | 4.8% | 4.2% |
| **Panic Disorder** | 2.4% | 3.1% |
| **Other\*\*** | 4.0% | 3.1% |

In addition, in order to better understand whether differences between this sample (summarized in Supplementary Table 1) and the sample used in Dinga et al. (*44*) may have influenced their power to detect statistically significant RSFC-clinical symptom correlations, we conducted supplementary analyses in a separate sample of N = 184 subjects (acquired during on-going studies at Cornell and Toronto) that more closely resembles their dataset. As in Dinga et al., this sample was acquired from four scanners (see below for details) and was a more diagnostically heterogeneous group: subjects were psychiatric outpatients presenting with symptoms of depression or anxiety and meeting DSM-IV criteria for a diagnosis of an anxiety disorder (26.1% of subjects, with generalized anxiety disorder, post-traumatic stress disorder, or panic disorder), major depressive disorder (53.2%), or both (20.7%), but there were no requirements for treatment resistance, and as in Dinga et al., they were not required to meet full DSM-IV criteria for a currently active major depressive episode (i.e. all MDD patients in this sample had experienced a major depressive episode, but some were in partial or full remission). Their mean age was 45.1 years, and 60.8% were female. Their medication status was as follows: 62.0% on an antidepressant; 12.0% on a mood stabilizer; 8.7% on an antipsychotic; and 53.8% taking another medication, including benzodiazepines, non-benzodiazepine sedative-hypnotics, thyroid hormone, and/or stimulants. Their mean HAMD total score was 17.5 with a range of 3 to 36.

All subjects provided written informed consent. All recruitment procedures and study protocols were reviewed and approved by the Institutional Review Boards of Weill Cornell Medicine or Toronto Western Hospital.

### MRI Data Acquisition

All subjects in the primary sample received a high-resolution T1-weighted anatomical MRI scan (MP-RAGE) and a T2*-weighted resting state functional MRI (gradient echo spiral in-out sequence) on a General Electric Signa 3T scanner. Scanning parameters for the Cornell subset were as follows: TR = 2 s, volumes = 180, FOV = 240 mm, slices = 28, XY resolution = 3.75 mm, Z resolution = 5 mm. Scanning parameters for the Toronto subset were as follows: TR = 2 s, volumes = 300, FOV = 220 mm, slices = 32, XY resolution = 3.44 mm, Z resolution = 5 mm.

As noted above, additional supplementary analyses were conducted in a second sample that more closely resembled the sample used in Dinga et al, and as in that report, subjects in this sample were scanned on one of four different scanners. The scanning parameters for two of the scanners (N = 25 and N = 98, respectively) were identical to those above. The third scanner (N = 19) used the following parameters: Siemens Trio 3T, TR = 2 s, volumes = 170, FOV = 224 mm, slices = 22, XY resolution = 3.5 mm, Z resolution = 5 mm. And the fourth scanner (N = 42) used the following parameters: Siemens Trio 3T, TR = 2.25 s, volumes = 166, FOV = 220 mm, slices = 39, XY resolution = 2.5 mm, Z resolution = 3.5 mm.

### fMRI Data Preprocessing and Functional Connectivity Quantification

Preprocessing was identical to the procedure defined in our previous report. To summarize the most important points, preprocessing of the resting state fMRI scans was implemented in AFNI and included standard procedures for slice-timing correction, spatial smoothing (4-mm FWHM Gaussian kernel), nonlinear registration to Montreal Neurological Institute (MNI) common space (via AFNI’s 3dQwarp function), temporal bandpass filtering (0.01–0.1 Hz), detrending (linear and quadratic), and regression on nuisance signals derived from head motion (12 parameters), CSF, white matter, and ANATICOR correction for local and global hardware artifacts. High-motion volumes (framewise displacement > 0.3 mm) were censored, as were the volumes preceding and following these high-motion frames. Importantly, nuisance signal regression and bandpass filtering were performed simultaneously in a single step (excluding subjects (8.9%) if the number of data points remaining after censoring was insufficient for performing this regression), as previous studies indicate that bandpass filtering prior to nuisance signal regression and censoring may lead to noise from high-motion volumes contaminating additional volumes (*45, 46*). The resulting residual time series volumes, registered to MNI common space, were used for functional connectivity quantification. See (*18*) for additional details, including validation of this protocol for controlling for motion-related artifacts.

#### Functional Connectivity Quantification

Again, this procedure was identical to the one specified in our previous work. Briefly, to reduce the dimensionality of the resulting rsfMRI residual time series data, we applied an established and extensively validated functional parcellation system delineating 258 functional network nodes (10-mm diameter spheres) spanning cerebral cortex, subcortical structures and cerebellum (*47*). These 258 nodes included 245 regions specified in the original parcellation by Power et al. (excluding 19 nodes from that parcellation that had incomplete MRI volume coverage or poor signal [SNR < 100] in our sample), plus 13 additional regions that have been implicated in depression with MNI coordinates (X, Y, Z mm) as follows: bilateral nucleus accumbens (+/−12, +8, −8); bilateral subgenual anterior cingulate (+/−4, +15, −11); bilateral caudate nucleus (+/−12, +18, −3); bilateral amygdala (+/−19, −2, −21); bilateral ventral hippocampus (+/−27, −15, −18); locus coeruleus (+/−3, −33, −27); ventral tegmental area (+/−6, −15, −15); and dorsal raphe nucleus (+/−6, −27, −21). BOLD signal time series were extracted from each ROI (averaging across all voxels), and a correlation matrix quantifying functional connectivity between each ROI and every other ROI was calculated for each subject using AFNI’s 3dNetCorr function. To aid in subsequent statistical analyses, we applied the Fisher *z*-transformation to these Pearson correlation coefficient matrices. We refer to these Fisher *z*-transformed correlation coefficients as resting state functional connectivity (RSFC) measures in all subsequent analyses.

#### Data Quality Assessment

In addition to the controls for motion-related artifacts described above, we also controlled for subject- and scanner-related differences in the temporal signal-to-noise ratio (SNR, defined as the mean of the MR signal over time / the standard deviation of the time series) by 1) excluding any ROI if its SNR was less than 100 in more than 5% of subjects; 2) including only voxels with SNR > 100 in calculating the mean BOLD signal time series for each ROI; and 3) excluding subjects with SNR < 100 in any of the remaining 258 nodes described above. Finally, we used multiple linear regression to further control for scanner and age effects by regressing the RSFC features (correlation coefficients) on subjects’ ages and dummy variables for site.

#### Preprocessing for Supplementary Analyses

As noted above, we performed supplementary analyses in a second sample, in order to better understand whether preprocessing and/or clinical sample differences between the analyses reported in Drysdale et al. (*18*) and Dinga et al. (*44*) may have influenced their power to detect statistically significant RSFC-clinical symptom correlations. For these supplementary analyses, we replicated the preprocessing stream described in Dinga et al. (*44*) and summarized in Supplementary Table 1. Although similar overall, key differences in this preprocessing stream included motion artifact correction via ICA-AROMA instead of motion-related nuisance signal regression; no censoring of high-motion volumes; no correction for scanner- or age-related differences; and no evaluation or correction for individual or scanner-related differences in SNR.

### Data Analysis

The stability and significance of correlations between HAMD clinical features and RSFC features was assessed by calculating the 33,123 Pearson correlation coefficients (PCCs) between each RSFC feature and one of the 16 HAMD item-level measures repeatedly on 1000 bootstrap replicates (each replicate consisted of n = 220 draws with replacement of matched RSFC and HAMD measures from the same subject). The 17th HAMD measure was excluded as it frequently had zero variance on bootstrap replicates. We used PCCs rather than Spearman correlations, as Spearman correlations like those used in (*18*) showed no noticeable improvement relative to PCCs for this analysis.

Following the procedure of Efron and colleagues (*48*), for each bootstrap replicate we converted the 33,123 PCCs between RSFC features and the chosen HAMD measure to *z*-values by taking the inverse cumulative distribution function (CDF) of the normal distribution applied to the *p*-values associated with each correlation *t*-statistic (see equation (1.2) and corresponding example in (*48*)). We then used estimates of the root-mean-square of the correlations between all pairs of *z*-values to correct the significance cutoff for the *z*-values to account for correlations among the RSFC features using the procedure detailed in (*48*). Applying this over the 1000 bootstrap replicates and taking the 2.5% and 97.5% percentiles of this empirical distribution (the “percentile bootstrap”; (*49*)) of the variance of the significant fraction of *z*-values cutoff variance (calculated for each replicate using equation 1.4 of (*48*)) led to the confidence intervals for the significant fractions shown in Fig. 1C for all 16 HAMD measures (yellow whiskers). All confidence intervals were corrected for the 16 separate tests using the Bonferroni-Holm step-down procedure (*50*) to adjust the percentiles used, and thus the 95% (corrected) confidence intervals shown are corrected for both correlations between RSFC features (using the procedure in (*48*)) and for multiple tests across HAMD features (using Holm-Bonferroni for 16 tests). We note that the Holm-Bonferroni correction is conservative, and the former led to only a modest correction, thus these results may be somewhat conservatively corrected overall. The bars shown in Fig. 1C were calculated using the mean observed significant area (past the two-tailed *p* < 0.05 significance threshold) observed across the 1000 bootstrap replicates of *z*-values, with the 2.5% of *z*-values expected to be above the two-tailed *p* < 0.05 threshold for the standard normal distribution subtracted so that chance level corresponds to zero. Confidence intervals reported for Fig 1. D were also calculated using the percentile bootstrap on the 1000 bootstrap replicates.

Canonical correlation analysis (*51, 52*) was performed between clinical measures and a selected subset of screened RSFC features (those with highest Spearman correlation) as previously described (*18*). Due to this feature screening step, we use validation on held-out data in subsequent analyses to avoid overly optimistic correlation estimates due to training-set overfitting (note that if we were specifically concerned with out-of-sample prediction, we should also have an additional fully out-of-sample test set, as recommended in the statistical learning (*53*) and neuroimaging (*54*) literature), but here we limit ourselves to a simple training and validation/test set to best focus on the stability of various approaches given limited data). To assess the stability of canonical variates resulting from CCA, 90% of matched RSFC/HAMD data was randomly sampled without replacement and used as a training set on which non-regularized or regularized CCA was fit, and the coefficients of this fit were then applied to the remaining 10% of test data to estimate the projection of these data into the CCA space as well as the resulting correlation between canonical variates on this held out validation data (this was done a total of 1000 times).

To better stabilize CCA coefficients, L2-regularized canonical correlation analysis (CCA), as described in (*55*) was also applied. This approach uses two regularization parameters, *λ_X_* and *λ_Y_* to regularize the estimated covariance matrices for the RSFC and clinical features, respectively. To find the best combination of these two variables, a grid search over possible values of the parameters was conducted, with 1000 RCCA fits found for each parameter combination (over *λ_X_* ∈ {0,0.1,1,10,100,1000, 1*e*6, 1*e*9} and *λ_Y_* ∈ {0,0.1,1,10,100, 1000, 1*e*6, 1*e*9}) as well as number of features selected chosen in 10 feature increments from 10 to 200 RSFC features. For each set of parameters, model fitting was done on training data and then assessed via the magnitude of the 1st canonical correlation coefficient on held out validation “test” data, using the same procedure as described above for standard CCA.

Analyses were conducted using a combination of custom Python and R code as well as R code for RCCA by (*55*), the Python Seaborn package (Michael Wascom) based on Matplotlib (*56*) for plotting, Jupyter Notebook, Ipython (*57*), Scipy (*58*) and Numpy (*59*).

## Results

### Testing for Robust Correlations between RSFCs and Clinical Symptoms

We begin with a modern approach to a classical problem: establishing the existence and strength of correlations between brain and behavior using mass univariate statistics. Examining number, strength, and effect size of these correlations gives us a strong basis from which to begin more complicated multivariate analyses, and convinces us of the utility of doing so. Further, understanding the structure of univariate correlations between RSFC and clinical symptoms gives us insight into what kind of challenges might present themselves in the multivariate setting.

To test the significance of some correlations among a large number of them, it is not uncommon to transform the *t*-statistics associated with each univariate correlation to *z*-values, which can then be compared against the standard normal distribution with mean 0 and variance 1 to test for significance. In the current application, we have 33,123 Pearson Correlation Coefficients (hereafter, PCCs), each of which can be converted into a corresponding *z*-value. However, these statistics are not independent of one another. The average correlation (±SD) between the RSFC features is 0.120 ± 0.0173, or only about 2–3 times less than the largest correlations between the RSFC features and HAMD clinical features. As it has been established that treating correlated *z*-values as though they are independent can significantly distort results using the distribution of *z*-values (*60*), we must correct for this. Conveniently, recent work on large-scale statistical estimation in similar scenarios provides us with tools to correct our estimates for just such correlations (*48*). Further, if we are to compare PCCs against all of 16 HAMD clinical measures we are interested in (we exclude HAMD 17 due to insufficient variance in this study), we will also have to take these multiple comparisons into account when assessing how many *z*-values are significant. To do this we use the Holm-Bonferroni correction to control family-wise error (*50*).

With these corrections in place, we can compare our distributions of *z*-values for correlations between RSFC and clinical features. We are interested in the number of *z*-values that exceeds the threshold of significance expected by chance (corrected for large-scale correlations and multiple comparisons; see Methods). We are also interested in establishing what the variance of the number of significant features is: is it stable, or do small changes in the data collection conditions translate to large changes in the number of correlations that are found to be significant (indicating unstable correlation estimates)? To estimate the variance of the number of correlations above the significance threshold, we use the bootstrap (*49*), resampling the RSFC and clinical data for each subject with replacement to generate 1000 bootstrap replicate data sets, and then run the *z*-value procedure from (*48*) on each replicate. The result of such a procedure for HAMD item 1 (HAMD1) is shown in Fig. 1A, where the histogram is of the median *z*-value for each RSFC to HAMD1 correlation over 1000 bootstrap replicates. The standard normal distribution against which we are comparing for significance (dotted red line) may be compared with a smoothed estimate of the median *z*-value distribution (green curve) and the excess area between these curves above threshold (black shading and arrow, vertical dotted lines are s ± 1.96) used as a measure of the expected number of *z*-values above the number expected by chance (where chance is 2.5% for *a* = 0.05 and a two-sided test).

Using the procedure in (*48*) to estimate the variance in the number of *z*-values above threshold for each bootstrap replicate *z*-value distribution, and then the “percentile bootstrap” (*49*) over our 1000 replicates (the 2.5% and 97.5% percentiles of the bootstrap distribution, for two-tailed *α* = 0.05), we can generate confidence intervals for the significant *z*-value estimates. These confidence intervals can then be corrected for the 16 multiple comparisons across the HAMD clinical features using the Holm-Bonferroni procedure, yielding the results shown in Fig. 1B. Here, each bar represents the estimated number of *z*-values found above chance level (2.5% for two-tailed *α* = 0.05) for all 16 HAMD measures considered, and the yellow whiskers show the 95% confidence interval for each (corrected for multiple comparisons). We can see that a number of the *z*-value distributions for the correlations have significantly more correlations than expected by chance. In particular, HAMD measures 1, 3, 5, 6, 7, 12, and 13 have median significant percentages (amount above 2.5%) well in excess of 1%, and overall, 14 out of the 16 *z*-value distributions show reliable, significant shifts in correlations. This pattern is recapitulated by the effect sizes (Fig. 1C) as measured by Cohen’s d (calculated between the smoothed *z*-value distributions like the green line in Fig. 1A with the standard normal), where *d* = 0.2 to *d* = 0.5 are considered small to medium effect sizes and *d* = 0.5 to *d* = 0.8 are considered medium to large effect sizes by convention (*61*).

Using the bootstrap replicates, we can also examine the range of RSFC to HAMD measure correlations. Fig. 1D shows the 1000 most positive (left) and 1000 most negative (right) correlations, ordered by the average PCC across bootstrap replicates (solid blue line). Superimposed we see the 95% confidence interval (percentile bootstrap), showing the range containing 95% of the PCC estimates over the 1000 bootstrap replicates. First, it is clear from these results that all of the most positive 1000 PCCs and a substantial fraction of the most negative 1000 PCCs are significant (their confidence interval excludes zero). This is what we would expect in general for the tails of a normal distribution, assuming correlations are relatively stable. The significant shift in the distribution we see in Fig. 1A can be seen in the upward shift of the positive correlations indicated by the black arrow and dotted line (Fig. 1D). We also note, however, that there is a significant range over which different bootstrap replicates might yield different orderings of the coefficients within, for example, the top 1000. We can see this even more clearly in Fig. 1E, which shows violin plots detailing the distribution of PCCs for the 10 most positive PCCs on average. Again these distributions are all significantly different from zero, with mean 95% confidence intervals ± SD of 0.148 ± 0.0152 to 0.376 ± 0.0119, but we note these distributions all look relatively exchangeable. That is, within the top 10 PCCs, the distributions look very similar and their ranking within the top 10 would change considerably over different bootstrap replicates. As described in more detail below, this is an important point when we consider feature selection using correlations, where a large number of very similar variables could result in highly variable feature selection.

### RSFC-clinical symptom correlations are sensitive to clinical sampling and preprocessing decisions

In a recent preprint, Dinga and colleagues reported the results of an analysis similar to our earlier work (*18*) and concluded that correlations between functional connectivity features and clinical symptoms were not statistically significant (*44*), which would seem to contradict the findings reported in Fig. 1. However, there were several important differences between these two studies, especially in their clinical sample characteristics and preprocessing pipelines (see Supplementary Table 1 for details). Of note, the sample in Ref. (*44*) included N = 87 subjects scanned on four different scanners (versus N=220 subjects scanned on just two scanners in our previous work, yielding a larger number of subjects per scanner and potentially more stable corrections for scanner-related differences). Dinga et al. did not directly control for scanner-related differences, and they took a different approach to controlling for motion and other global signal artifacts. Importantly, their sample was also more clinically heterogeneous (including MDD, generalized anxiety disorder, social phobia, or panic disorder with no specified requirements for active depressive symptoms vs. currently active, treatment-resistant MDD in our work). By testing for RSFC-clinical symptom correlations in this more heterogeneous sample, the approach in Ref. (*44*) assumes that the mechanisms that drive these correlations are the same across all these disorders, but this may not be true. For example, it is possible that different mechanisms may drive anxiety symptoms in MDD compared panic disorder, in which case an analysis of subjects with mixed diagnoses could yield smaller effect sizes and unstable results in held-out data.

To test whether these clinical sample and preprocessing differences could influence their power to detect robust RSFC-clinical symptom correlations, we repeated the analysis reported in Fig. 1 in a second more clinically heterogeneous sample of ISM84 subjects with MDD or an anxiety disorder, scanned on one of four scanners, and preprocessed exactly as in Ref. (*44*). (See Methods and Materials for additional details.) The results are depicted in Supplementary Fig. 1. They show that statistically significant RSFC-clinical symptom correlations are still detectable for 10 of 16 symptoms (vs. 14 of 16 in Fig. 1B), but these associations are modest compared to those in Fig. 1B, with uniformly small effect sizes (d = 0.21–0.29 for 5 symptoms, d < 0.2 for all others). These results are consistent with the interpretation that distinct mechanisms give rise to RSFC-clinical symptom correlations across these heterogeneous disorders and that preprocessing decisions could also be important.

### Stable canonical correlations between RSFC features and clinical symptoms

As noted above, Fig. 1D indicates that the ranking of the top 10 PCCs could change considerably over different bootstrap replicates. This in turn suggests that a large number of very similar variables could result in highly variable feature selection across bootstrap replicates. Such redundant structure in the correlations motivates searching for low-rank embedding of the data that might find a small number of dimensions in which to express the many redundant correlations between RSFC features and clinical measures. Canonical Correlation Analysis (CCA) (*51, 52*) is a classical multi-view statistical approach that in this context can be used to find linear combinations of RSFC measures and clinical features that have maximum correlation with each other. Each linear combination defines a set of canonical variates (CVs), one for the RSFC measures and one for the clinical measures, that define a two-dimensional embedding of the data in which the data are maximally correlated. After the first CVs are found, subsequent CVs orthogonal to the previous ones can be estimated, defining a low-rank embedding of both the RSFC and clinical data into a linear subspace with a rank equal to the number of CVs estimated (*55*). Thus, if effective, CCA might provide a low-dimensional representation of the relationship between neuroimaging and clinical features in the form of a simplified summary of the interesting structure between the brain and clinical symptoms, as well as the ability to generalize to new data, and potential targets for causal investigations.

However, traditional CCA has some potential weaknesses, particularly on large-scale, correlated data. In particular, previous literature has shown that CCA coefficients become unstable in the presence of multicollinearity (significant correlations between variables, as we might suspect between both the RSFC features) (*55*). Further, CCA can only operate on as many variables as there are observations, so that feature selection is necessary prior to applying CCA in order to reduce the 33,123 potential RSFC measures to a number less than or equal to the number of observations (the number of subjects in the study)(*55*). Despite this, CCA has yielded promising results on the data presented in our previous work and re-analyzed here, although as direct assessment of the stability of CCA solutions was not integral to the other analyses in that study (*18*), it was left to subsequent work.

Here, we resample the data 1000 times (without replacement) into training (90% of subjects) and validation (“test”) sets (the remaining 10% of subjects) in order to assess the stability of CCA on this data across different numbers of features selected (see Methods for details). The results can be seen in Fig. 2A, where we show that standard CCA does seem to overfit, as the training correlations in the 1st CV subspace gradually approach values above 0.9 (although they break down as the number of features approaches the number of subjects, which is 198 for 90% of the data), while the test canonical correlations increase initially (peaking at 20 features) but then decrease towards 0.1 by 190 features. The violin plots also show that variance of the distributions for test canonical correlations is quite large, with a significant fraction of the distribution making excursions below zero canonical correlation as the number of selected features included increases. Still, the best fit has a median canonical correlation of 0.557 (with 1st and 3rd quartile range across replicates of [0.456, 0.642]), suggesting the approach is promising.

We hypothesized that these results might be stabilized via L2-regularization applied to the CCA coefficients associated with both the RSFC and clinical features, as both were multicollinear. We also suspected that the RSFC coefficients in particular would benefit from regularization as the number of features included increased. Such regularization is common in the high-dimensional regression literature (where it is frequently referred to as the “ridge” penalty (*62*)) and induces a small downward bias in coefficient magnitude in exchange for what in practice is often a large reduction in coefficient variance (*53*). In regularized CCA (RCCA), we shrink both the sample covariance matrix for the RSFC features 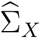 and for the clinical measures 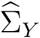 toward the identity matrix by replacing them with 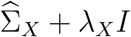 and 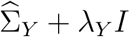, respectively (55). This requires specifying the value of the two regularization parameters *λ_X_* and *λ_Y_* for each RCCA fit. To assess the effects of these parameters on fit quality, we fit each of our RCCA models over a grid of *λ_X_* and *λ_Y_*, with each parameter taking values in set {0, 0.1, 1, 10, 100, 1000, 1*e*6, 1*e*9}.

The median canonical correlation results on the held-out test data (over 1000 replicates) are shown in Fig. 2B. The left panel shows one grid (for 160 selected features) in detail, and we can see that just a small amount of regularization of the RSFC feature coefficients greatly improves the test canonical correlations, as an increase of *λ_X_* from 0 to 0.01 increases test canonical correlations across all values of *λ_Y_*. While somewhat less sensitive to changes in *λ_Y_*, regularization of the HAMD coefficients also benefits the fit, and we see a peak median test canonical correlation at *λ_X_* = 0.1 and *λ_Y_* = 1.0 of 0.735 (with 1st and 3rd quartile range of [0.665, 0.797] across replicates). The right panel of Fig. 2B shows grids for the other number of features shown (over the same grid of regularization parameters). We can also visualize the distributions of the training and test 1st canonical correlations as we did in Fig. 2A, but now for the best RCCA fit at *λ_X_* = 0.1, *λ_Y_* = 1.0. We see that compared to the CCA fit in Fig. 2A, the test canonical correlation distributions have lower variance, remain well above zero, and appear to improve with increasing number of features. We can visualize the difference between the CCA and best RCCA directly by overlaying the median and 1st through 3rd quartiles for the test canonical correlation derived from taking the 25% through 75% percentiles of the 1000 train/test subsample replicates over both approaches (Fig. 2D; RCCA in red, CCA in gray). These show clearly that CCA test canonical correlations (on the 1st CV) steadily approaches zero as more RSFC features are included, while RCCA only improves up to 160 features and after that reliably maintains high canonical correlations across replicates (out to the maximum number we tried here: 200 features). Further, if we examine the stability of test correlations between additional canonical variates (Fig. 2E), we see that RCCA (taken at its best performance at 160 RSFC features with *λ_X_* = 0.1, *λ_Y_* = 1.0) uniformly outperforms CCA (taken at its best performance at 20 RSFC features) for the first 4 sets of canonical variates. Thus we find that L2-regularization of both RSFC feature and HAMD measure coefficients stabilizes and improves low-dimensional co-embedding of neuroimaging and clinical measures.

It is also interesting to consider further the number of screened features accommodated by the best RCCA fit versus the best CCA fit (i.e., what selecting 160 features looks like compared to selecting 20). Fig. 2F and G show, over the 1000 subsamples used as the training data, the ranked distributions of which RSFC features were chosen by the screening procedure (*18*). It is clear that if we look at how frequently the most selected features were included across replicates, having just 20 features (Fig. 2F) means just 3 features are selected more than 80% of the time, whereas having 160 features results in 25 features appearing more than 80% of the time. If we run pairwise comparisons looking at how many features appear in both of two replicates (randomly choosing 100 of the subsample replicates) we see that the number of overlapping features selected by the feature selection increases linearly with the total number of features selected (Fig 2H). Thus, stabilizing CCA with regularization here allows the model to leverage more features than standard CCA, yielding a broader set of more reliable features to be used in the resulting low-dimensional representation that result in higher test correlations.

## Discussion

Together, these results show that linear combinations of RSFC features are stable and statistically significant predictors of distinct clinical symptom combinations in MDD patients who are actively depressed. As shown in our previous work (*18*), individual depressed patients, in turn, can be clustered into subgroups defined by relatively homogeneous patterns of altered functional connectivity in depression-related brain networks, which predict distinct clinical symptom profiles. Of course, it remains unclear whether and how RSFC alterations in specific circuits are driving depression-related symptoms and behaviors, or merely correlated with them. Optogenetic tools, which can be integrated with fMRI and other noninvasive neuroimaging modalities, offer one approach to addressing this question.

First introduced in 2010 (*43*), this method combines high-field fMRI with opsins to manipulate the activity of genetically defined cellular subtypes and test for both local and global effects on neuronal activity and brain network function. This initial report by Lee et al. (*43*) underscored two of the most important and commonly implemented applications of optogenetic fMRI (ofMRI). First, it showed how ofMRI could be used to glean mechanistic insights into the neurophysiological basis of the fMRI BOLD signal—a critical issue for interpreting the results of clinical neuroimaging studies. This report (*43*) showed that optogenetic stimulation of neocortical or thalamic excitatory neurons was sufficient to drive local BOLD signal responses, informing an ongoing debate about the nature of the neuronal signals that underlie the BOLD signal, as well as the cellular subtypes that give rise to them. Subsequent ofMRI studies showed that the BOLD signal is more strongly correlated with local spiking activity than with the local field potential (*63*) and is driven by the effects of neuronal activity on cerebral venules (*64*).

Second, Lee et al. (*43*) went on to show how ofMRI could be used for whole-brain functional circuit mapping, by optogenetically manipulating the activity of excitatory pyramidal cells in a specific brain area and testing for downstream BOLD signal effects. More recent studies have extended this approach to map and differentiate the functional networks activated by specific circuits (e.g. dorsal vs. ventral hippocampus, and other hippocampal subregions)(*65–69*) and by specific cellular subtypes (e.g. striatal medium spiny neurons expressing D1 vs. D2 dopamine receptors, and dopaminergic vs. glutamatergic cells in the ventral tegmental area)(*70–72*), often with surprising results that could not be predicted based solely on mapping the axonal projection fields of a given brain region (e.g. Ref. (*69*)).

Of particular relevance for translational neuroscience studies, of MRI methods can also be easily extended to recapitulate disease-related pathophysiological processes, evaluate their impact on brain-wide functional network dynamics, and test for causal effects on behavior. To this end, we illustrate one approach for formulating hypotheses regarding subtype-specific mechanisms driving depressive symptoms and behaviors, and testing them in animal models using optogenetic fMRI (Fig. 3A), drawing on the results of two recently published works. In this model, resting state fMRI (or potentially other neuroimaging modalities) can be used to identify functional connectivity features in candidate circuits that predict depression-related symptoms and behaviors. Optogenetic fMRI, in turn, can be used to recapitulate and validate these connectivity changes in functionally related circuits in rodents, and test for causal effects on specific depression-related behaviors. One approach to identifying promising candidate circuits involves searching for connectivity alterations and clinical symptoms that tend to co-occur. For example, our previous clinical neuroimaging work used CCA to define a low-dimensional representation of connectivity features in specific circuits that predicted specific symptom combinations (*18*). Hierarchical clustering on the first two canonical variates revealed at least four clusters or subtypes (Fig. 3B) predicting group differences in multiple symptoms, especially anhedonia and anxiety (Fig. 3C). Group differences in anhedonia and anxiety, in turn, were associated with group differences in RSFC in depression-related brain networks (Fig. 3D).

These subtype-specific patterns were complex, involving hundreds of connectivity alterations in dozens of neuroanatomical areas. However, qualitatively, two observations stood out. First, Subtypes 1 and 4 were associated with increased anxiety and deficits in functional connectivity in fronto-amygdala circuits (see green, dashed-line boxes in Fig. 3D), which have been implicated in the regulation of fear memories and the cognitive reappraisal of negative emotional states (*73–76*). Second, Subtypes 3 and 4 were associated with increased anhedonia and elevated functional connectivity between the medial prefrontal cortex, ventral striatum, and other frontostriatal circuits that have been implicated in reward processing, effort valuation, and motivation (*6, 23, 77–83*).

Optogenetic tools provide one means of testing whether altering functional connectivity in these circuits is sufficient for driving specific depression-related behaviors. Stable step function opsins (SSFOs) are particularly useful in this context, in that they were designed to achieve stable, partial depolarization in genetically defined cell types on a timescale of minutes (*30*), suitable for use in resting state fMRI analyses of low-frequency signal fluctuations, but still immediately reversible, enabling within-subject statistical comparisons. Furthermore, by partially depolarizing neurons and rendering them more excitable and responsive to their physiological inputs, they can in principle be used to reversibly modulate functional connectivity in specific circuits and cell types.

At least one recent optogenetic fMRI study by Ferenczi et al. (*36*) provides evidence consistent with the hypothesis that increased functional connectivity in a specific frontostriatal network, qualitatively similar to the pattern observed in Subtypes 3 and 4, is sufficient to drive anhedonic behavior in rats. In this study, SSFO was expressed in CaMKIIa+ projection neurons in the medial prefrontal cortex (mPFC), and rsfMRI was used to quantify functional connectivity changes elicited by SSFO activation in the mPFC (Fig. 3E). SSFO activation increased functional connectivity between the mPFC target and a network of structures including the ventral striatum, nucleus accumbens, orbitofrontal cortex, anterior cingulate cortex, and thalamus (Fig. 3E), qualitatively similar to many of the areas exhibiting increased functional connectivity in Subtypes 3 and 4. SSFO modulation of mPFC projection cells was also sufficient to drive anhedonia-like behavior in the sucrose preference test (Fig. 3F), and individual differences in functional connectivity in a frontostriatal circuit were correlated with sucrose preference behavior (Fig. 3G).

Importantly, this approach also provides a means of testing how circuits interact to produce anhedonic behavior. Ferenczi et al. (*36*) went on to show that mPFC and the ventral tegmental area (VTA) compete to influence processing in striatum (data not shown). VTA stimulation drove a striatal BOLD response that predicted reward-seeking behavior, while SSFO modulation of mPFC excitability suppressed the striatal response to VTA stimulation and disrupted reward processing. Of course, these findings do not necessarily indicate that the same mechanism is involved in driving anhedonic behavior in Subtypes 3 and 4. Rather, they show that this particular pattern of frontostriatal hyperconnectivity, elicited by increasing the excitability of mPFC projection neurons, is sufficient to disrupt reward-seeking behavior. Future studies could test whether these subtypes are associated with hyperexcitability in mPFC; with deficits in striatal reward reactivity; and with abnormal interactions between VTA, mPFC, and striatum. Likewise, new viral tools for targeting opsin expression to topologically defined projection neuron subtypes with increased ease and efficiency (*84–86*) will enable more targeted investigations that modulate connectivity between specific nodes in this frontostriatal network.

The example in Fig. 3 illustrates one approach to formulating hypotheses about candidate circuits driving subtype-specific dysfunction in depression, based on qualitatively similar connectivity alterations that tend to co-occur with specific symptoms or behaviors across subtypes. However, it is worth noting that candidate circuits could also be identified in a data-driven way, especially with larger sample sizes. Indeed, multi-view methods like RCCA are well suited to this purpose, in that they reduce a large number of complex correlations at the brain and clinical symptom level to a low dimensional space that captures the most significant relationships between them in a concise and orthogonal way (yielding particular combinations of RSFCs and clinical symptoms that covary, orthogonal to others). Reliable coefficients in the RCCA model suggest targets for bidirectional optogenetic control in rodent experiments that could be used to test if symptom dimensions can indeed be dissociated by modulating the candidate neural targets.

It is also worth noting some important limitations and caveats associated with this approach. First, Fig. 2A underscores how CCA has a tendency to overfit when combined with a feature selection step. Therefore, when screening is used to pre-select features for further analysis, careful training and test validation are necessary to generate models that perform well in held-out data and to avoid overly optimistic results due to overfitting. Second, our approach to feature selection is adequate for identifying stable and robust associations between RSFC features and clinical symptoms, but other methods could yield superior results. In particular, future studies will likely use nonlinear multi-view and/or sparse methods or other more advanced feature selection protocols to improve the results reported here, although the general results and conceptual framework will likely be similar.

Third, these approaches may be highly sensitive to clinical sample characteristics (e.g. distinct circuit mechanisms may be at play in active depression, depression in remission, and various anxiety disorders), as well as to data quality, head motion, and other sources of global signal artifacts. Therefore, it is important to optimize and validate preprocessing methods and other data quality controls, based on the goals of a given study. Finally, categorical subtyping is just one approach to parsing diagnostic heterogeneity, and the 4-cluster solution in Fig. 3B is not the only solution. Rather, as discussed in more detail in (*18*), this 4-cluster solution was stable and clinically useful (predicting clinical symptoms and treatment response), but also most likely constrained by features of the subtype discovery dataset in (*18*), especially sample size. Likewise, although a model anchored in categorical subtypes provides a familiar and clinically useful heuristic for clinicians to parse diagnostic heterogeneity in depression, other methods might be superior. One alternative approach that warrants further examination would substitute a multi-dimensional rating system for categorical subtype diagnoses.

These caveats notwithstanding, the results in Figs. 1–3 and the accompanying review highlight the potential for integrating clinical neuroimaging analyses with optogenetic fMRI studies to formulate and test hypotheses regarding subtype-specific mechanisms driving particular symptoms and behaviors in depression. RCCA can be used to discover robust and stable associations between functional connectivity and behavior, linking specific circuits with specific clinical symptom combinations that may be differentially involved in individual MDD patients. Optogenetic fMRI, in turn, provides a powerful tool for testing hypotheses derived from clinical neuroimaging data; for implicating specific patterns of network dysfunction as causal mechanisms, not just functional correlates; and for isolating the contributions of specific network nodes and circuits and studying their interactions.

## Acknowledgments

The authors gratefully acknowledge Dr. Karl Deisseroth and Dr. Emily Ferenczi for granting permission to adapt selected figure panels from Ref. (*36*), for presentation in Fig. 3. This work was supported by grants from the National Institute of Mental Health, the One Mind Institute, the Klingenstein-Simons Foundation Fund, the Rita Allen Foundation, the Whitehall Foundation, the Dana Foundation, the Brain and Behavior Research Foundation (NARSAD), and the Hartwell Foundation. L.G. was supported by the Simons Foundation Society of Fellows. M.J.D. has received research grants from Neuronetics and Tal Medical, Inc. All other authors report no biomedical financial interests or other potential conflicts of interest.

---

**Supp. Table 1.**
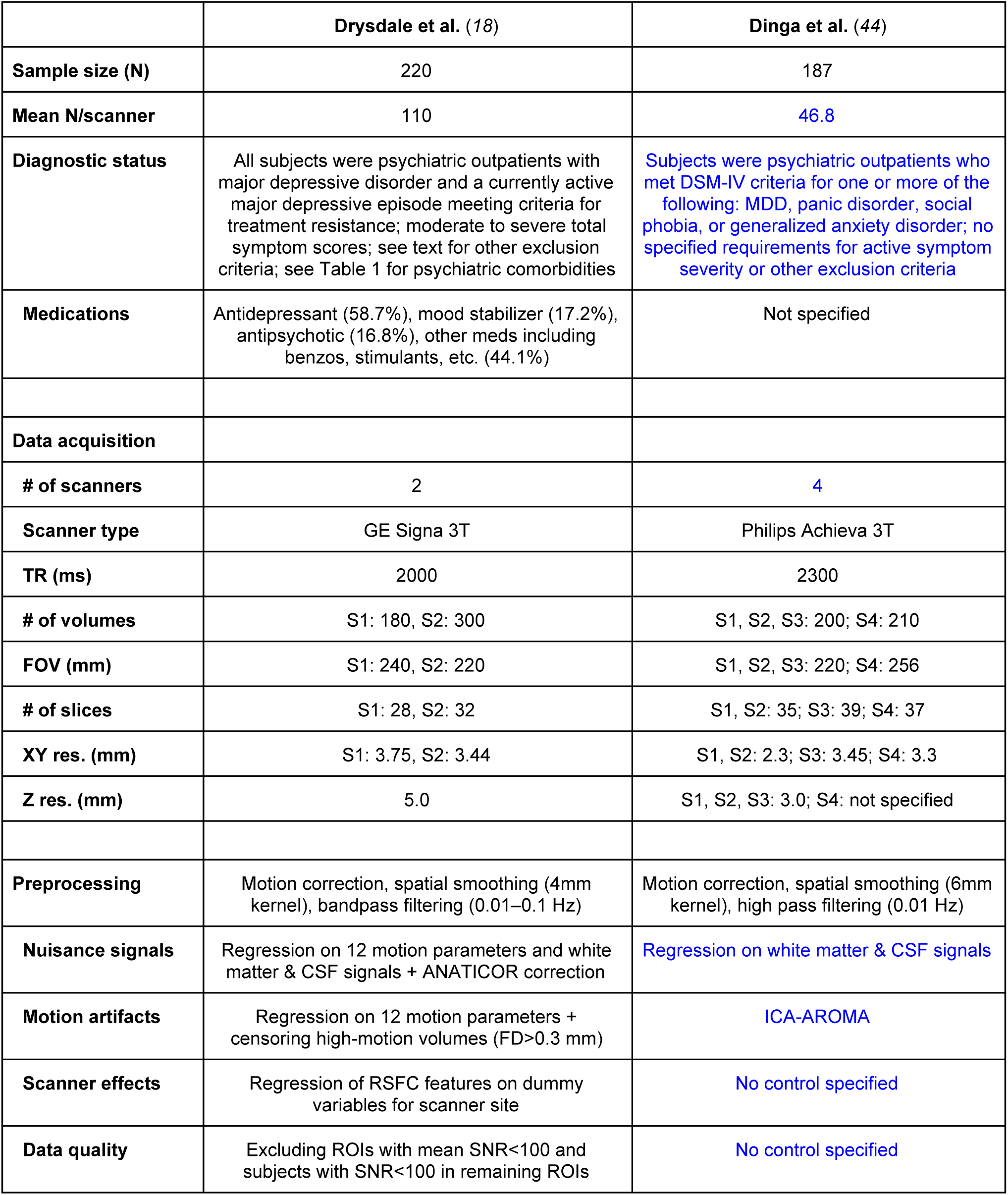
Contrasting sample characteristics and preprocessing in Refs. (*18*) and (*44*), with potentially important differences highlighted in blue. S# = scanner 1, 2, 3, and 4.

**Supplementary Figure 1:**
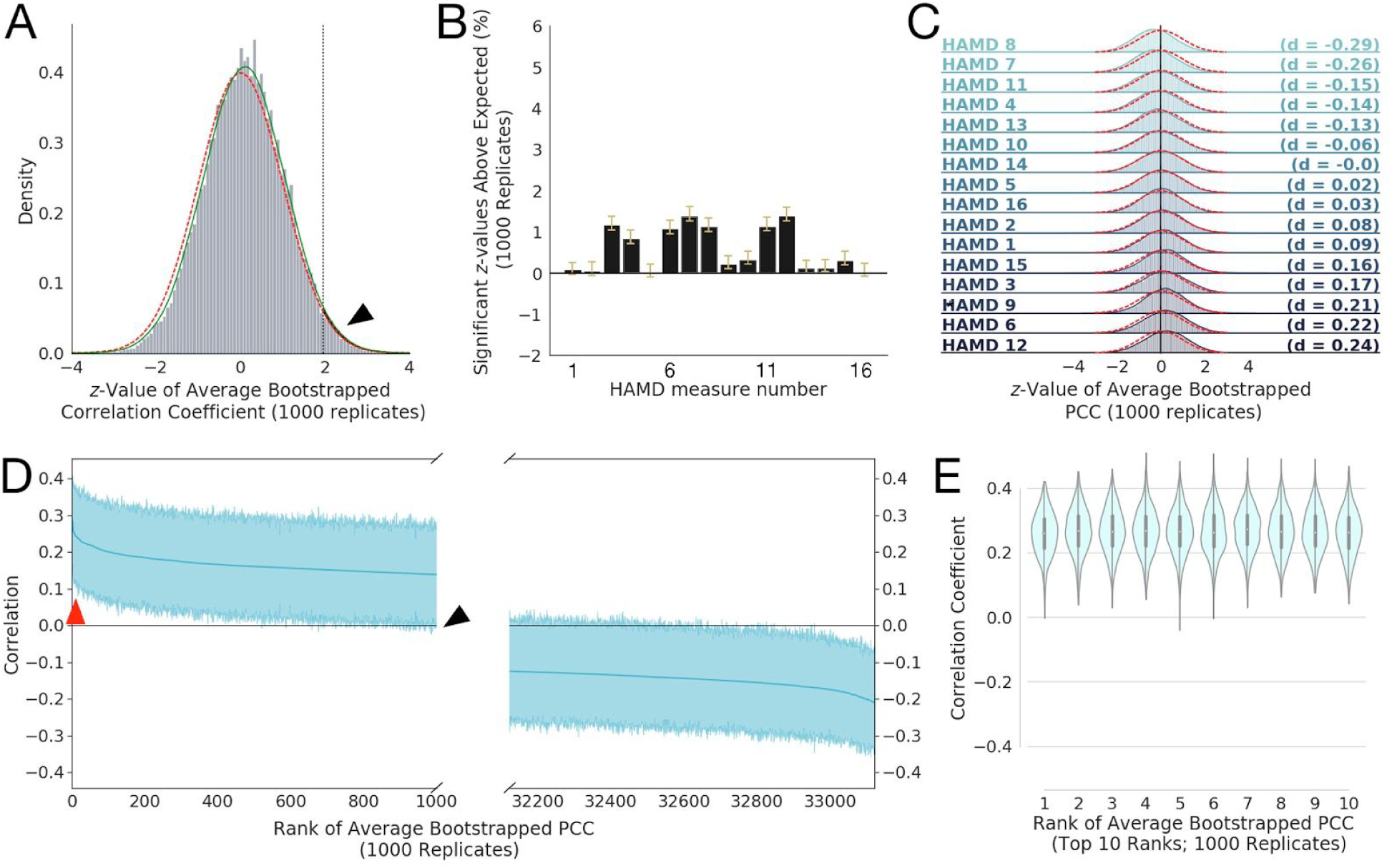
Modest RSFC-clinical symptom correlations in a heterogeneous multi-site sample (*44*). **A)** Histogram of *z*-values for the 33,123 average Pearson Correlation Coefficients (PCCs) between Resting-State Functional Connectivity (RSFC) features and HAMD clinical measures (averaged over 1000 bootstrap replicates), compared to a standard 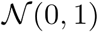 Gaussian distribution (red line) and with a smoothed kernel density estimate plotted over the histogram (green line; see Methods), as in Fig. 1A. **B)** Bar plots of the mean percentage of *z*-values that exceeded that expected by chance (e.g., the percentage above 2.5%, shown as the shaded black area in **A** for HAMD 1) for 1000 bootstrap replicates. Yellow whiskers on the bars denote 95% confidence intervals (corrected for multiple comparisons and data correlation using Bonferroni-Holm and (*48*), respectively). Comparing with Fig. 1B, we note a severe disruption of the correlation pattern. **C)** Histograms of *z*-values like that shown in **A** for all 16 HAMD clinical measures considered, ordered by effect size (Cohen’s d), given at right of each plot. Magnitudes between 0.2 and 0.5 are considered small to medium effect sizes, between 0.5 and 0.8 are considered medium to large effect sizes. Here, none exceed 0.3 in magnitude). Red dotted lines denote the standard normal distribution. Asterisk (*) marks the distribution for HAMD1 shown in **A**. Again, compared with Fig 1C, we see a marked decrease in effected sizes. **D)** Bootstrapped PCCs for HAMD measure 1 for the most positive 1000 (left) and 1000 most negative (right) RSFC features (shaded regions shows 95% percentile-bootstrap confidence interval for the mean), ordered by mean correlation (thick central blue line). Red arrow points to top 10 most positive-ranked RSFC features (shown in **E)** and the black arrow to where we would expect an upwardly significant difference as in Fig. 1D, but now see none. **E)** Violin plot (with superimposed boxplots showing 1st and 3rd quartiles as black bar and the median as white point) of the top ten positive ranked RSFCs by average PCC to HAMD measure 1 (corresponding to red arrow in **D**), with mean 95% confidence intervals ± SD of [0.132 ± 0.0197, 0.391 ± 0.0145], Coefficient variability appears similar to that in Fig. 1.

## References

1. T. Insel et al., Research Domain Criteria (RDoC): Toward a New Classification Framework for Research on Mental Disorders. Am. J. Psychiatry. 167, 748–751 (2010).

2. T. R. Insel, B. N. Cuthbert, Brain disorders? Precisely. Science. 348, 499–500 (2015).

3. D. J. Oathes, B. Patenaude, A. F. Schatzberg, A. Etkin, Neurobiological Signatures of Anxiety and Depression in Resting-State Functional Magnetic Resonance Imaging. Biol. Psychiatry. 77, 385–393 (2015).

4. M. Goodkind et al., Identification of a Common Neurobiological Substrate for Mental Illness. JAMA Psychiatry. 2015/02/05 (2015), doi: 10.1001/jamapsychiatry.2014.2206.

5. R. J. Davidson, D. Pizzagalli, J. B. Nitschke, K. Putnam, Depression: Perspectives from affective neuroscience. Annu. Rev. Psychol. 53, 545–574 (2002).

6. D. A. Pizzagalli, in *Annual Review of Clinical Psychology, Vol 10*, T. D. Cannon, T. Widiger, Eds. (2014), vol. 10 of Annual Review of Clinical Psychology, pp. 393–423.

7. M. L. Wong et al., Pronounced and sustained central hypernoradrenergic function in major depression with melancholic features: relation to hypercortisolism and corticotropin-releasing hormone. Proc. Natl. Acad. Sci. U. S. A. 97, 325–330 (2000).

8. P. W. Gold, G. P. Chrousos, Organization of the stress system and its dysregulation in melancholic and atypical depression: high vs low CRH/NE states. Mol. Psychiatry. 7, 254–275 (2002).

9. B. J. Carroll et al., A SPECIFIC LABORATORY TEST FOR THE DIAGNOSIS OF MELANCHOLIA - STANDARDIZATION, VALIDATION, AND CLINICAL UTILITY. Arch. Gen. Psychiatry. 38, 15–22 (1981).

10. A. J. Lewy, R. L. Sack, L. S. Miller, T. M. Hoban, Antidepressant and Circadian Phase-Shifting Effects of Light. Science. 235, 352–354 (1987).

11. A. F. Schatzberg et al., Neuropsychological deficits in psychotic versus nonpsychotic major depression and no mental illness. Am. J. Psychiatry. 157, 1095–1100 (2000).

12. B. A. Clementz et al., Identification of Distinct Psychosis Biotypes Using Brain-Based Biomarkers. Am. J. Psychiatry. 173, 373–384 (2016).

13. S. K. Hill et al., Neuropsychological impairments in schizophrenia and psychotic bipolar disorder: findings from the Bipolar-Schizophrenia Network on Intermediate Phenotypes (B-SNIP) study. Am. J. Psychiatry. 170, 1275–1284 (2013).

14. R. E. Amir et al., Rett syndrome is caused by mutations in X-linked MECP2, encoding methyl-CpG-binding protein 2. Nat. Genet. 23, 185–188 (1999).

15. S. Baron-Cohen, S. Wheelwright, The empathy quotient: an investigation of adults with Asperger syndrome or high functioning autism, and normal sex differences. J. Autism Dev. Disord. 34, 163–175 (2004).

16. J. Sebat et al., Strong association of de novo copy number mutations with autism. Science. 316, 445–449 (2007).

17. D. Pinto et al., Functional impact of global rare copy number variation in autism spectrum disorders. Nature. 466, 368–372 (2010).

18. A. T. Drysdale et al., Resting-state connectivity biomarkers define neurophysiological subtypes of depression. Nat. Med. 23, 28–38 (2017).

19. M. D. Greicius et al., Resting-state functional connectivity in major depression: Abnormally increased contributions from subgenual cingulate cortex and thalamus. Biol. Psychiatry. 62, 429–437 (2007).

20. L. Pezawas et al., 5-HTTLPR polymorphism impacts human cingulate-amygdala interactions: a genetic susceptibility mechanism for depression. Nat. Neurosci. 8, 828–834 (2005).

21. Y. I. Sheline, J. L. Price, Z. Yan, M. A. Mintun, Resting-state functional MRI in depression unmasks increased connectivity between networks via the dorsal nexus. Proc. Natl. Acad. Sci. U. S. A. 107, 11020–11025 (2010).

22. Y. I. Sheline et al., The default mode network and self-referential processes in depression. Proc. Natl. Acad. Sci. U. S. A. 106, 1942–1947 (2009).

23. B. Knutson, J. P. Bhanji, R. E. Cooney, Atlas, Lauren Y., I. H. Gotlib, Neural responses to monetary incentives in major depression. Biol. Psychiatry. 63, 686–692 (2008).

24. C. Liston et al., Default Mode Network Mechanisms of Transcranial Magnetic Stimulation in Depression. Biol. Psychiatry. 76, 517–526 (2014).

25. S. J. Broyd et al., Default-mode brain dysfunction in mental disorders: a systematic review. Neurosci. Biobehav. Rev. 33, 279–296 (2009).

26. A. Anand et al., Activity and connectivity of brain mood regulating circuit in depression: a functional magnetic resonance study. Biol. Psychiatry. 57, 1079–1088 (2005).

27. E. S. Boyden, F. Zhang, E. Bamberg, G. Nagel, K. Deisseroth, Millisecond-timescale, genetically targeted optical control of neural activity. Nat. Neurosci. 8, 1263–1268 (2005).

28. J. A. Cardin et al., Driving fast-spiking cells induces gamma rhythm and controls sensory responses. Nature. 459, 663–667 (2009).

29. F. Zhang et al., Multimodal fast optical interrogation of neural circuitry. Nature. 446, 633–639 (2007).

30. O. Yizhar et al., Neocortical excitation/inhibition balance in information processing and social dysfunction. Nature. 477, 171–178 (2011).

31. B. Y. Chow et al., High-performance genetically targetable optical neural silencing by light-driven proton pumps. Nature. 463, 98–102 (2010).

32. L. Grosenick, J. H. Marshel, K. Deisseroth, Closed-loop and activity-guided optogenetic control. Neuron. 86, 106–139 (2015).

33. A. R. Adamantidis, F. Zhang, A. M. Aravanis, K. Deisseroth, L. de Lecea, Neural substrates of awakening probed with optogenetic control of hypocretin neurons. Nature. 450, 420–424 (2007).

34. V. Gradinaru, M. Mogri, K. R. Thompson, J. M. Henderson, K. Deisseroth, Optical deconstruction of parkinsonian neural circuitry. Science. 324, 354–359 (2009).

35. A. V. Kravitz et al., Regulation of parkinsonian motor behaviours by optogenetic control of basal ganglia circuitry. Nature. 466, 622–626 (2010).

36. E. A. Ferenczi et al., Prefrontal cortical regulation of brainwide circuit dynamics and reward-related behavior. Science. 351, aac9698 (2016).

37. H. E. Covington et al., Antidepressant Effect of Optogenetic Stimulation of the Medial Prefrontal Cortex. Journal of Neuroscience. 30, 16082–16090 (2010).

38. D. Chaudhury et al., Rapid regulation of depression-related behaviours by control of midbrain dopamine neurons. Nature. 493, 532–+ (2013).

39. K. M. Tye et al., Amygdala circuitry mediating reversible and bidirectional control of anxiety. Nature. 471, 358–362 (2011).

40. S. Lammel et al., Input-specific control of reward and aversion in the ventral tegmental area. Nature. 491, 212–217 (2012).

41. G. D. Stuber et al., Excitatory transmission from the amygdala to nucleus accumbens facilitates reward seeking. Nature. 475, 377–380 (2011).

42. S. Ciocchi et al., Encoding of conditioned fear in central amygdala inhibitory circuits. Nature. 468, 277–282 (2010).

43. J. H. Lee et al., Global and local fMRI signals driven by neurons defined optogenetically by type and wiring. Nature. 465, 788–792 (2010).

44. R. Dinga et al., Evaluating the evidence for biotypes of depression: attempted replication of Drysdale et.al. 2017. *bioRxiv* (2018), p. 416321.

45. J. Carp, Optimizing the order of operations for movement scrubbing: Comment on Power et al. Neuroimage. 76, 436–438 (2013).

46. J. D. Power, K. A. Barnes, A. Z. Snyder, B. L. Schlaggar, S. E. Petersen, Steps toward optimizing motion artifact removal in functional connectivity MRI; a reply to Carp. Neuroimage. 76, 439–441 (2013).

47. J. D. Power et al., Functional Network Organization of the Human Brain. Neuron. 72, 665–678 (2011).

48. B. Efron, Correlated *z*-values and the accuracy of large-scale statistical estimates. J. Am. Stat. Assoc. 105, 1042–1055 (2010).

49. B. Efron, R. J. Tibshirani, An Introduction to the Bootstrap (CRC Press, 1994).

50. S. Holm, A Simple Sequentially Rejective Multiple Test Procedure. Scand. Stat. Theory Appl. 6, 65–70 (1979).

51. H. Hotelling, Relations Between Two Sets of Variates. Biometrika. 28, 321–377 (1936).

52. D. R. Hardoon, S. Szedmak, J. Shawe-Taylor, Canonical correlation analysis: an overview with application to learning methods. Neural Comput. 16, 2639–2664 (2004).

53. T. Hastie, R. Tibshirani, J. Friedman, The Elements of Statistical Learning: Data Mining, Inference, and Prediction, Second *Edition* (Springer-Verlag New York, ed. 2, 2009), *Springer Series in Statistics*.

54. L. Grosenick, B. Klingenberg, K. Katovich, B. Knutson, J. E. Taylor, Interpretable whole-brain prediction analysis with GraphNet. Neuroimage. 72, 304–321 (2013).

55. I. Gonzalez, S. Dejean, CCA: An R Package to Extend Canonical Correlation Analysis (available at https://core.ac.uk/download/pdf/6303071.pdf).

56. J. D. Hunter, Matplotlib: A 2D Graphics Environment. Comput. Sci. Eng. 9, 90–95 (2007).

57. F. Perez, B. E. Granger, IPython: A System for Interactive Scientific Computing. Computing in Science Engineering. 9, 21–29 (2007).

58. T. E. Oliphant, SciPy: Open source scientific tools for Python. Computing in Science and Engineering. 9, 10–20 (2007).

59. T. E. Oliphant, A guide to NumPy (Trelgol Publishing USA, 2006), vol. 1.

60. A. B. Owen, Variance of the number of false discoveries. J. R. Stat. Soc. Series B Stat. Methodol. 67, 411–426 (2005).

61. J. Cohen, Statistical power analysis for the behavioral sciences 2nd edn (1988).

62. A. E. Hoerl, R. W. Kennard, Ridge Regression: Biased Estimation for Nonorthogonal Problems. Technometrics. 12, 55–67 (1970).

63. I. Kahn et al., Optogenetic drive of neocortical pyramidal neurons generates fMRI signals that are correlated with spiking activity. Brain Res. 1511, 33–45 (2013).

64. X. Yu et al., Sensory and optogenetically driven single-vessel fMRI. Nat. Methods. 13, 337–340 (2016).

65. Z. Liang et al., Mapping the functional network of medial prefrontal cortex by combining optogenetics and fMRI in awake rats. Neuroimage. 117, 114–123 (2015).

66. R. W. Chan et al., Low-frequency hippocampal-cortical activity drives brain-wide resting-state functional MRI connectivity. Proc. Natl. Acad. Sci. U. S. A. 114, E6972–E6981 (2017).

67. A. T. L. Leong et al., Long-range projections coordinate distributed brain-wide neural activity with a specific spatiotemporal profile. Proc. Natl. Acad. Sci. U. S. A. 113, E8306–E8315 (2016).

68. A. J. Weitz et al., Optogenetic fMRI reveals distinct, frequency-dependent networks recruited by dorsal and intermediate hippocampus stimulations. Neuroimage. 107, 229–241 (2015).

69. N. Takata et al., Optogenetic Activation of CA1 Pyramidal Neurons at the Dorsal and Ventral Hippocampus Evokes Distinct Brain-Wide Responses Revealed by Mouse fMRI. PLOS One. 10, e0121417 (2015).

70. D. Bernal-Casas, H. J. Lee, A. J. Weitz, J. H. Lee, Studying Brain Circuit Function with Dynamic Causal Modeling for Optogenetic fMRI. Neuron. 93, 522–532.e5 (2017).

71. H. J. Lee et al., Activation of Direct and Indirect Pathway Medium Spiny Neurons Drives Distinct Brain-wide Responses. Neuron. 91, 412–424 (2016).

72. M. Brocka et al., Contributions of dopaminergic and non-dopaminergic neurons to VTA-stimulation induced neurovascular responses in brain reward circuits. Neuroimage. 177, 88–97 (2018).

73. T. D. Wager, M. L. Davidson, B. L. Hughes, M. A. Lindquist, K. N. Ochsner, Prefrontal-subcortical pathways mediating successful emotion regulation. Neuron. 59, 1037–1050 (2008).

74. M. R. Milad, G. J. Quirk, Neurons in medial prefrontal cortex signal memory for fear extinction. Nature. 420, 70–74 (2002).

75. E. A. Phelps, M. R. Delgado, K. I. Nearing, J. E. LeDoux, Extinction learning in humans: role of the amygdala and vmPFC. Neuron. 43, 897–905 (2004).

76. K. N. Ochsner, J. J. Gross, The cognitive control of emotion. Trends Cogn. Sci. 9, 242–249 (2005).

77. D. A. Pizzagalli et al., Reduced Caudate and Nucleus Accumbens Response to Rewards in Unmedicated Individuals With Major Depressive Disorder. Am. J. Psychiatry. 166, 702–710 (2009).

78. D. Kvitsiani et al., Distinct behavioural and network correlates of two interneuron types in prefrontal cortex. Nature. 498, 363–366 (2013).

79. K. L. Hillman, D. K. Bilkey, Neural encoding of competitive effort in the anterior cingulate cortex. Nat. Neurosci. 15, 1290–1297 (2012).

80. P. L. Croxson, M. E. Walton, J. X. O’Reilly, T. E. J. Behrens, M. F. S. Rushworth, Effort-based cost-benefit valuation and the human brain. J. Neurosci. 29, 4531–4541 (2009).

81. M. T. Treadway, N. A. Bossaller, R. C. Shelton, D. H. Zald, Effort-based decision-making in major depressive disorder: a translational model of motivational anhedonia. J. Abnorm. Psychol. 121, 553–558 (2012).

82. W. Schultz, P. Dayan, P. R. Montague, A neural substrate of prediction and reward. Science. 275, 1593–1599 (1997).

83. R. N. Cardinal, J. A. Parkinson, J. Hall, B. J. Everitt, Emotion and motivation: the role of the amygdala, ventral striatum, and prefrontal cortex. Neurosci. Biobehav. Rev. 26, 321–352 (2002).

84. D. G. R. Tervo et al., A Designer AAV Variant Permits Efficient Retrograde Access to Projection Neurons. Neuron. 92, 372–382 (2016).

85. L. Luo, E. M. Callaway, K. Svoboda, Genetic Dissection of Neural Circuits: A Decade of Progress. Neuron. 98, 256–281 (2018).

86. K. T. Beier et al., Circuit Architecture of VTA Dopamine Neurons Revealed by Systematic Input-Output Mapping. Cell. 162, 622–634 (2015).

